# Structural defects caused by Acrodermatitis Enteropathica mutations in the extracellular domain account for mistrafficking and malfunction of ZIP4

**DOI:** 10.1101/2020.08.16.253294

**Authors:** Eziz Kuliyev, Chi Zhang, Dexin Sui, Jian Hu

## Abstract

ZIP4 is a representative member of the Zrt-/Irt-like protein (ZIP) transporter family and responsible for zinc uptake from diet. Loss-of-function mutations of human ZIP4 (hZIP4) drastically reduce zinc absorption, causing a life-threatening autosomal recessive disorder, Acrodermatitis Enteropathica (AE). Although the zinc transport machinery is located in the transmembrane domain conserved in the entire ZIP family, half of the missense mutations occur in the extracellular domain (ECD) of hZIP4, which is only present in a fraction of mammalian ZIPs. How the AE-causing mutations in the ECD lead to ZIP4 malfunction has not be fully clarified. In this work, we characterized all the seven confirmed AE-causing missense mutations in hZIP4-ECD and found that the variants exhibited completely abolished zinc transport activity measured in a cell-based transport assay. Although the variants were able to be expressed in HEK293T cells, they failed to traffic to cell surface and were largely retained in the ER with immature glycosylation. When the corresponding mutations were introduced in the ECD of ZIP4 from Pteropus Alecto, a close homolog of hZIP4, the variants exhibited impaired protein folding and reduced thermal stability, which likely account for intracellular mistrafficking of the AE-associated variants and as such a total loss of zinc uptake in cells. This work provides a molecular pathogenic mechanism for AE, which lays out a basis for potential therapy using small molecular chaperones.

## Introduction

Zinc is an essential micronutrient for any living organism. As the second most abundant transition metal element in humans after iron, zinc is required for function of over 2,000 transcription factors and activity of approximately 300 enzymes (1). It has been estimated that zinc binds to nearly 3,000 proteins which account for ~10% of the proteins encoded in human genome (2). Studies have shown that zinc deficiency may affect multiple biological processes, causing growth retardation, immune dysfunction, diarrhea, delayed sexual maturation, and skin lesions (3). Zinc deficiency is usually caused by inadequate zinc supply in foods or acquired diseases either reducing zinc uptake or increasing zinc loss (4). In rare cases, inherited zinc deficiency, which is an autosomal recessive disorder called Acrodermatitis Enteropathica (AE), is caused by loss-of-function (LOF) mutations of ZIP4 (5,6), the exclusive high-affinity zinc transporter in gastrointestinal system responsible for zinc uptake from regular diet (7–10). AE is fatal without treatment, but lifelong high-dose zinc supplementation on daily basis can effectively alleviate the symptoms (11), implying the presence of additional low-affinity zinc absorption mechanism(s).

ZIP4 is a representative member of the Zrt-/Irt-like Protein (ZIP) family (Solute Carrier 39A, SLC39A) consisting of divalent transition metal transporters ubiquitous in all the kingdoms of life (12–18). ZIP4 is specifically expressed on the apical side of enterocytes in small intestine and also in kidney where it is believed to be involved in zinc reabsorption from urine (18). Topologically, ZIP4 has a transmembrane domain (TMD) which is generally conserved in the entire ZIP family and a large extracellular domain (ECD) which is only present in a fraction of the family members (19,20). Be of interest, the AE-causing mutations are evenly distributed along the 12 exons of the zip4 gene without showing hotspot (21,22). As a result, half of the missense AE-causing mutations are mapped in the ECD and the other half in the TMD where the zinc transport machinery is located. A previous work has investigated several corresponding mutations in the TMD of mouse ZIP4 (mZIP4) but only one mutation in the ECD (equivalent to P200L in human ZIP4, hZIP4) was studied in the same report (7). The P200L variant of hZIP4 was characterized in a recent report (23). So far, it is still unclear about how the other AE-causing mutations in the ECD affect hZIP4 function and the corresponding molecular mechanisms.

The crystal structure of the ECD of ZIP4 from Pteropus Alecto (black fruit bat, pZIP4-ECD), which shares 68% sequence identity with hZIP4-ECD, provides a structural framework to deduce structural impacts of the AE-causing mutations on this regulatory domain required for optimal zinc transport (19). In this work, we functionally characterized all the seven confirmed AE-causing mutations in a human cell line (HEK293T) and biophysically studied the purified variants of pZIP4-ECD harboring the corresponding mutations. We found that all these mutations showed little zinc transport activity in a cell-based transport assay, although their overall expression levels were not much different from that of the wild type protein. The variants exhibited drastically reduced cell surface level and were aberrantly retained in the ER with immature glycosylation. For the purified pZIP4-ECD variants, biophysical studies revealed that the mutations caused reduced α-helical content, undermined packing and decreased thermal stability. Taken together, these findings indicate that impaired protein folding and compromised structural compactness caused by the AE-causing mutations in ZIP4-ECD account for ZIP4 dysfunction linked with intracellular mistrafficking.

## Results

### The AE-causing mutations in the ECD of hZIP4 led to total loss of zinc transport activity

A total of 17 missense mutations in hZIP4-ECD have been documented, of which seven are confirmed AE-causing mutations and the others are benign single-nucleotide polymorphisms (SNPs), according to the MASTERMIND database (https://mastermind.genomenon.com/) and an early collection (21). Mapping these residues on the hZIP4-ECD structural model (generated using SWISS-MODEL (24)) shows that the residues subjected to AE-causing mutations (C62R, R95C, A99T, N106K, P200L, Q303H and C309Y) are all in the structured regions, whereas most of the residues where the SNPs occur are on loops or in disordered regions (Figure 1). In this work, we characterized all the seven AE-causing mutations, including four homozygous mutations (C62R, P200L, Q303H and C309Y) (5,6,25,26) and three heterozygous mutations (R95C, A99T and N106K) (6,26,27), and studied their impacts on hZIP4 function. The hZIP4 variants with a C-terminal HA tag were transiently expressed in HEK293T cells and applied to the cell-based radioactive zinc transport assay (6,7,28,29). As shown in Figure 2A, in sharp contrast to the wild type hZIP4, none of the cells expressing any variants uptook more zinc than the blank group in which the cells were transfected with an empty vector, which means that the variants have no detectable zinc transport activity with 10 µM of Zn^2+^ added in culture media. For the wild type hZIP4, the previous studies have shown that zinc transport activity reaches plateau at 10 µM of Zn^2+^ under the same condition (28,29). To examine whether any activity can be detected at higher zinc concentration, the P200L variant was tested at various zinc concentrations up to 50 µM, but the results only confirmed no detectable activity (Figure S1).

**Figure 1.**
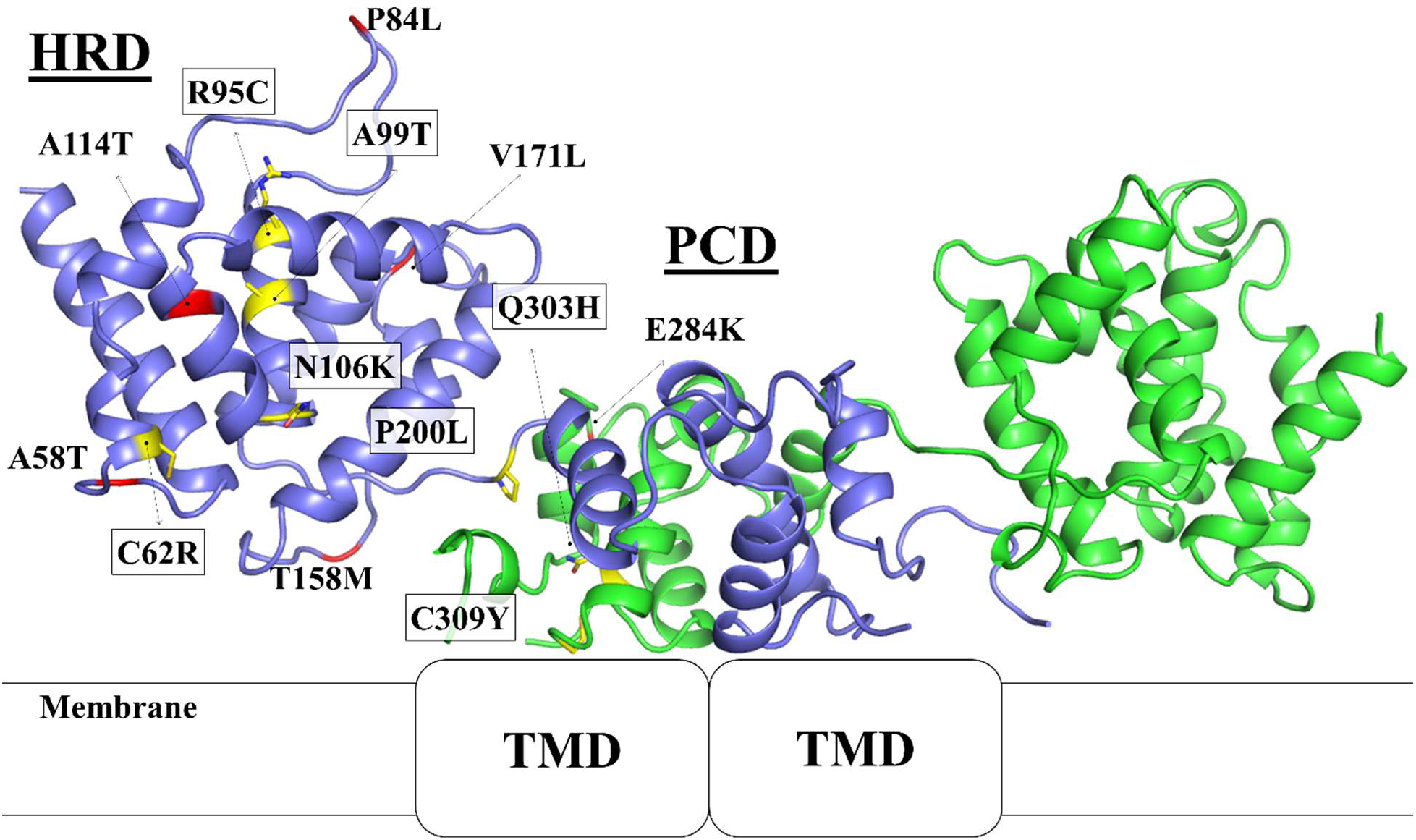
Mapping of the AE-causing mutations and the SNPs on the structural model of hZIP4-ECD dimer. Seven residues subjected to AE-causing mutations (highlighted in boxes) are in yellow and stick mode with four in the HRD domain and three in the PCD domain, whereas the residues at which SNPs occur are in red. Additional 3 SNPs not shown in the model are either in the highly disordered his-rich loop (R251W) or in the signal peptide (V2A and E10A).

**Figure 2.**
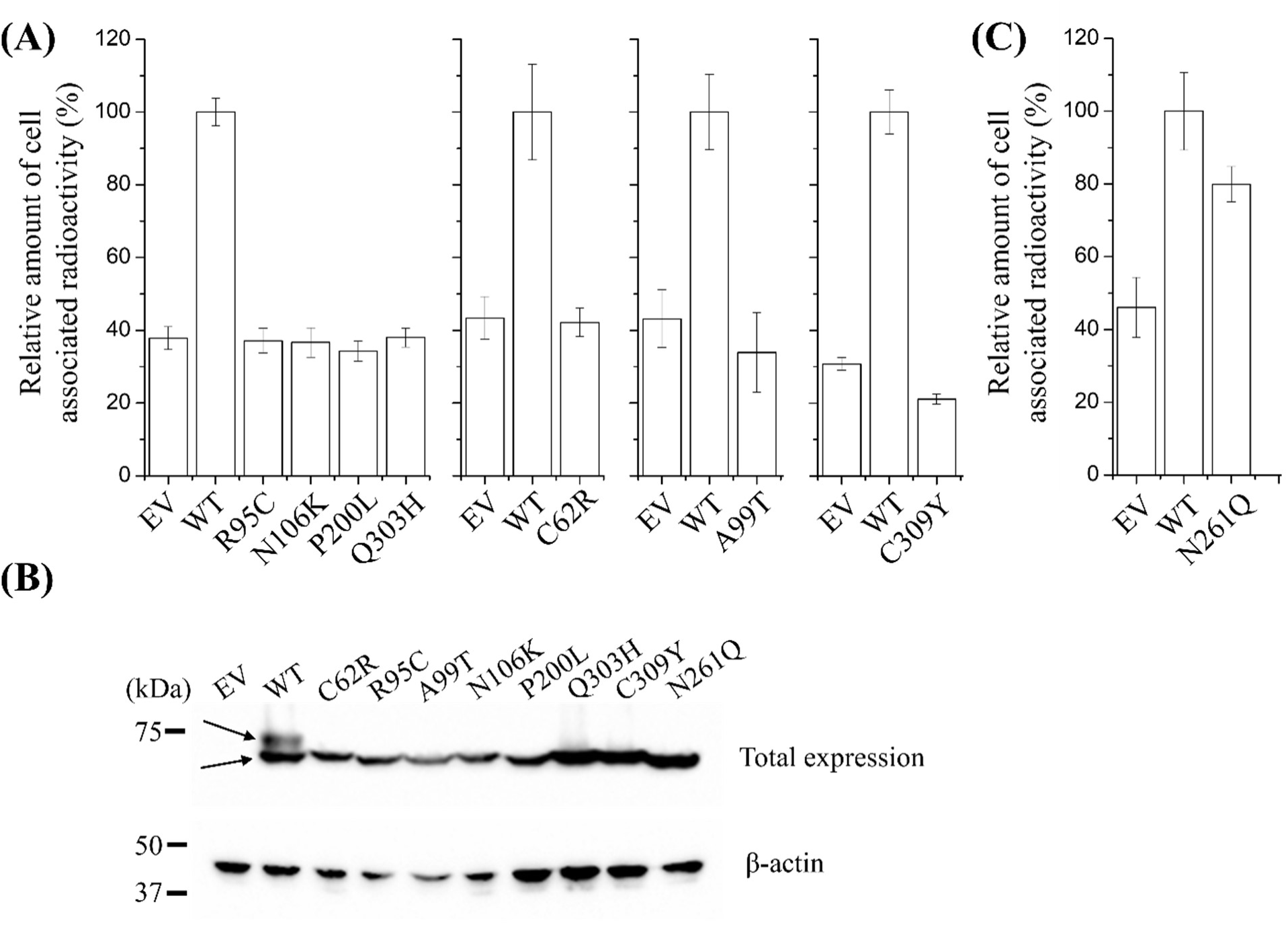
Functional characterization of hZIP4 and the variants. (A) Cell-based zinc transport assay of the AE-associated variants with 10 µM Zn^2+^ in the medium. The results of one out of 2-6 independent experiments are shown. Three technical repeats were included in each experiment. The error bars indicate ±1 standard deviation. (B) Total expression hZIP4 and the variants detected by Western blot using anti-HA antibody. The upper arrow indicates the maturely glycosylated hZIP4, whereas the lower arrow indicates the immaturely glycosylated form. β-actin was detected in the same sample as loading control. (C) Zinc transport assay of the N261Q variant. Three technical repeats were included in the experiment. The error bars indicate ±1 standard deviation.

### The AE-associated variants exhibited immature glycosylation

The previous study on mZIP4 has shown that the AE-causing mutations in the TMD led to significantly lowered expression levels, likely due to accelerated degradation (7). To examine whether the AE-causing mutations in the ECD have similar effects, the expression levels of the variants were detected in the whole cell samples using Western blots with an anti-HA antibody. As shown in Figure 2B, the overall expression levels of the variants are comparable to or modestly lower than that of the wild type hZIP4. However, the wild type hZIP4 has two discrete bands at approximately 75 kDa, whereas the AE-associated variants have only one corresponding to the lower band of the wild type protein. It has been determined that the upper band represents the maturely glycosylated form whereas the protein in the lower band is immaturely glycosylated (7), which is further confirmed in this work using PNGase F, an enzyme cleaving N-linked glycoside bond(s) (Figure S2). Thus, the total expression of any variants is not affected by the mutations in the ECD, but a defect in glycosylation was observed for all the AE-associated variants.

### N-glycosylation is not required for hZIP4 activity

Proteins expressed at cell surface are often modified by N-glycosylation, which may play a key role in protein function (30). As the AE-associated variants losing zinc transport activity is concomitant with glycosylation defects, we asked whether the lack in glycosylation is responsible for activity loss. Using the NetNGlyc server (http://www.cbs.dtu.dk/services/NetNGlyc/), we identified a potential glycosylation site at N261 in an “NxS/T” motif. We then generated the N261Q variant and expressed it in HEK293T cells. As expected, the N261Q variant showed only the lower band in Western blot (Figure 2B), confirming N261 is indeed the only N-glycosylation site in hZIP4. Importantly, zinc transport assay of the N261Q variant only showed a modest reduction of transport activity (Figure 2C), indicating that glycosylation at N261 is not pivotal for zinc transport and therefore lack of glycosylation of the AE-associated variants does not account for loss of zinc transport activity.

### The AE-associated variants have significantly diminished cell surface level

As the glycan are added to the client membrane proteins in the ER and processed in the ER and then in the Golgi in a stepwise manner (31), immature glycosylation is an indicator of defect in intracellular trafficking. To examine potential mislocalization, we tested cell surface expression levels of the AE-associated variants by applying the anti-HA antibody to the non-permeabilized cells fixed with formaldehyde. After extensive wash, non-specific bound antibody was removed, leaving those specifically bound with the C-terminal HA tag at cell surface. As shown in Figure 3A, all the AE-associated variants had substantially reduced cell surface levels when compared to the wild type hZIP4 in Western blots. Consistent with the zinc transport data, the N261Q variant missing the N-linked glycan had significantly higher level of cell surface expression than the variants linked with disease, indicating that the *N*-glycosylation at N261 is not a key factor for hZIP4 surface expression. We then applied immunofluorescence imaging to locate the AE-associated variants expressed in cells. To do that, the cells transiently expressing the wild type or the variants were fixed and permeabilized with formaldehyde and TX-100, followed by staining with an FITC-labeled anti-HA antibody. The samples were then checked under confocal laser scanning microscope (CLSM). Consistent with the Western blot results shown in Figure 2B, the wild type and the variants were all expressed in HEK293T cells at comparable levels with most of the staining signals within the cytoplasm (Figure 3B). To detect the transporters expressed at cell surface, the cells were fixed with formaldehyde and treated with the same anti-HA antibody, followed by extensive washing and checking under CLSM. In Figure 3B, fluorescence signals at cell surface can be convincingly detected for the wild type hZIP4 and the N261Q variant, whereas the signals of the AE-associated variants imaged under the same conditions were much weaker or not detectable, which is consistent with the Western blot data shown in Figure 3A. Collectively, our data strongly indicate that the cell surface expression levels of the AE-associated variants are largely suppressed, which is likely responsible for loss of zinc transport activity.

**Figure 3.**
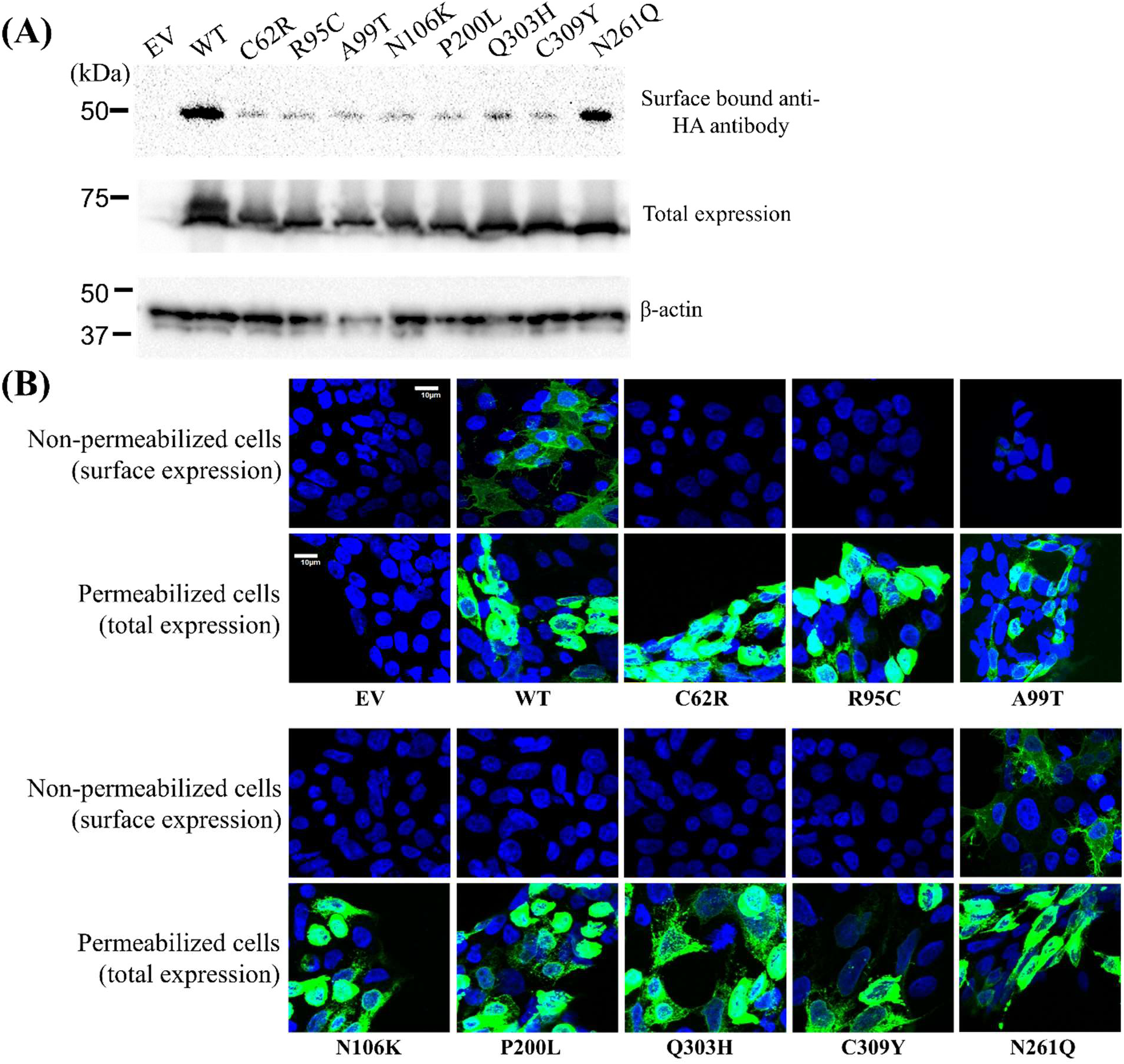
Cell surface expression of hZIP4 and its variants. (A) Surface expression detected by Anti-HA antibody in Western blot. Total expression of hZIP4 and the variants from the same batch of cells are shown for comparison. β-actin in the same sample was detected as loading control. (B) Detection of expression of hZIP4 and its variants. FITC-labeled anti-HA antibody was used to stain HA tagged hZIP4 in non-permeabilized cells (upper row, for detection of cell surface expression) or permeabilized cells (lower row, for detection of total expression in whole cells). The scale bar = 10 µm.

### The AE-associated variants are retained in the ER

Since all the AE-associated variants were expressed but not presented at cell surface, we then asked where the variants are located in cells. To determine intracellular location of the variants, we examined co-localization of hZIP4 and its variants with an ER-resident protein, calreticulin. As shown in Figure 4, the staining of the wild type hZIP4-HA (red) is partially overlapped with those of calreticulin (green), resulting in a yellow/orange color for the overlapped portion. This result suggests that a portion of hZIP4 in HEK293T cells are in the ER, at least partially due to overexpression driven by the strong CMV promoter in transient transfection. However, as significant amount of hZIP4 is not co-localized with calreticulin, this portion of the wild type hZIP4 is either in other organelles (such as the Golgi complex) or at the plasma membrane. In contrast, the AE-associated variants are mostly co-localized with calreticulin, strongly suggesting that the variants are primarily retained in the ER. As membrane proteins are initially glycosylated in the ER and then the glycans are further processed in the Golgi complex for maturation, the variants being retained in the ER is consistent with immature glycosylation of the variants. It is known that mistrafficking can be the result of protein misfolding and/or structural alteration imposed by disease-causing mutations, we are wondering whether and how the AE-causing mutations affect protein structure. Given the difficulty of obtaining adequate amount of purified full length hZIP4 or hZIP4-ECD to conduct biophysical characterization, we turned to the isolated pZIP4-ECD, which we have previously crystallized and biochemically characterized (19,28). Given the high sequence identity, we believe that the results obtained on purified pZIP4-ECD would help to understand how the disease-causing mutations affect the biophysical properties of the human counterpart.

**Figure 4.**
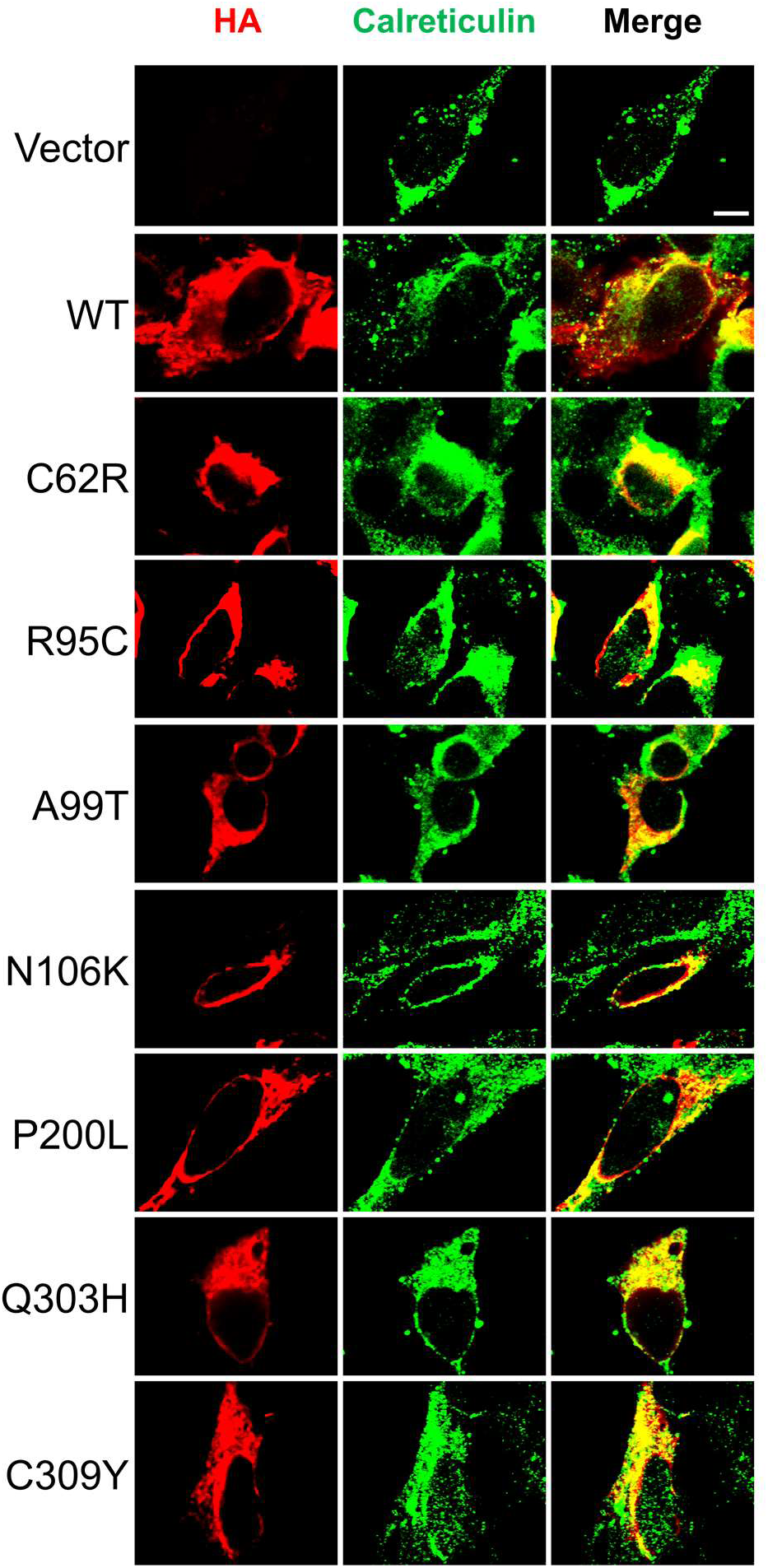
Colocalization of hZIP4 with ER marker calreticulin. Representative confocal images (100X) of HEK293T cells transiently expressing hZIP4-HA or the variants are shown. hZIP4 and its variants were detected with an anti-HA tag antibody and an Alexa-568 goat anti-mouse antibody (red). Calreticulin was detected with an anti-calreticulin antibody and an Alexa-488 anti-rabbit antibody (green). Scale bar = 5 µm.

### The AE-causing mutations alter secondary structure of pZIP4-ECD

We introduced mutations in pZIP4-ECD on the residues topologically equivalent to those in hZIP4 (C64R, R87C, A91T, D98K, P193L, Q299H and C305Y) and expressed the variants in the same way as for the wild type protein. We have noticed that, even for the wild type pZIP4-ECD, a large portion of purified protein from the Co-affinity column are not well folded as indicated in ion-exchange chromatography experiment, which is likely due to the presence of eight invariant cysteine residues required for four disulfide bonds formation. As a result, only a small fraction of expressed protein can be purified to homogeneity for later structural and biophysical experiments, making the overall yield of the wild type pZIP4-ECD barely higher than 1 mg per liter culture. For the variants, we found the misfolding issue is more severe, which led to further lower yield for most of the variants (0.1-0.5 mg/L). Particularly, the yields of two variants (D98K and C305Y) were too low to allow for further characterization. Therefore, we focused on the other five variants in later study.

During purification, we noticed that all the variants were eluted in size-exclusion chromatography with an apparent molecular weight of approximately 60 kDa, indicative of a homodimer as the wild type protein, so the AE-causing mutations do not affect ECD dimerization. The purified proteins were applied to circular dichroism (CD) spectrometer and the spectral data were analyzed on the K2D3 server to estimate secondary structure contents (32). As shown in Figure 5, the CD spectrum of the wild type pZIP4-ECD is of high α-helical content (estimated to be 66%) with the characteristic minima at 208 nm and 222 nm and maximum at 195 nm, which is consistent with the reported crystal structure (19). When compared to the wild type protein, all the variants have lower levels of helical content - C64R (61%), R87C (55%), A91T (58%), P193L (64%) and Q299H (56%). We also noticed the ratios of ellipticities (θ) at 222 nm and 208 nm were reduced for all the variants. The θ_222_/θ_208_ ratio is an indicator of whether the helices are packed to form coiled-coil like tertiary structures (33,34). When the value is smaller than 1, it may indicate poorly packed helices and lack of coiled-coils. For the wild type protein θ_222_/θ_208_ is 1.08, which is consistent with the crystal structure where two helix bundles form two separated subdomains: in the N-terminal helix-rich domain (HRD), α4 is surrounded and packed with the other eight helices, whereas four helices (α10-α12) in the C-terminal domain (PAL motif-containing domain, PCD) are packed against with the counterparts from the other monomer to form a homodimer (19). For all the tested variants, we found the θ_222_/θ_208_ ratios were reduced to be smaller than one - C64R (0.93), R87C (0.83), A91T (0.93), P193L (0.98) and Q299H (0.84). Therefore, lower helical content and reduced θ_222_/θ_208_ ratio revealed by the CD spectra indicate defects in protein folding for all the variants, of which R87C and Q299H have greatest effects whereas P193L has the least.

**Figure 5.**
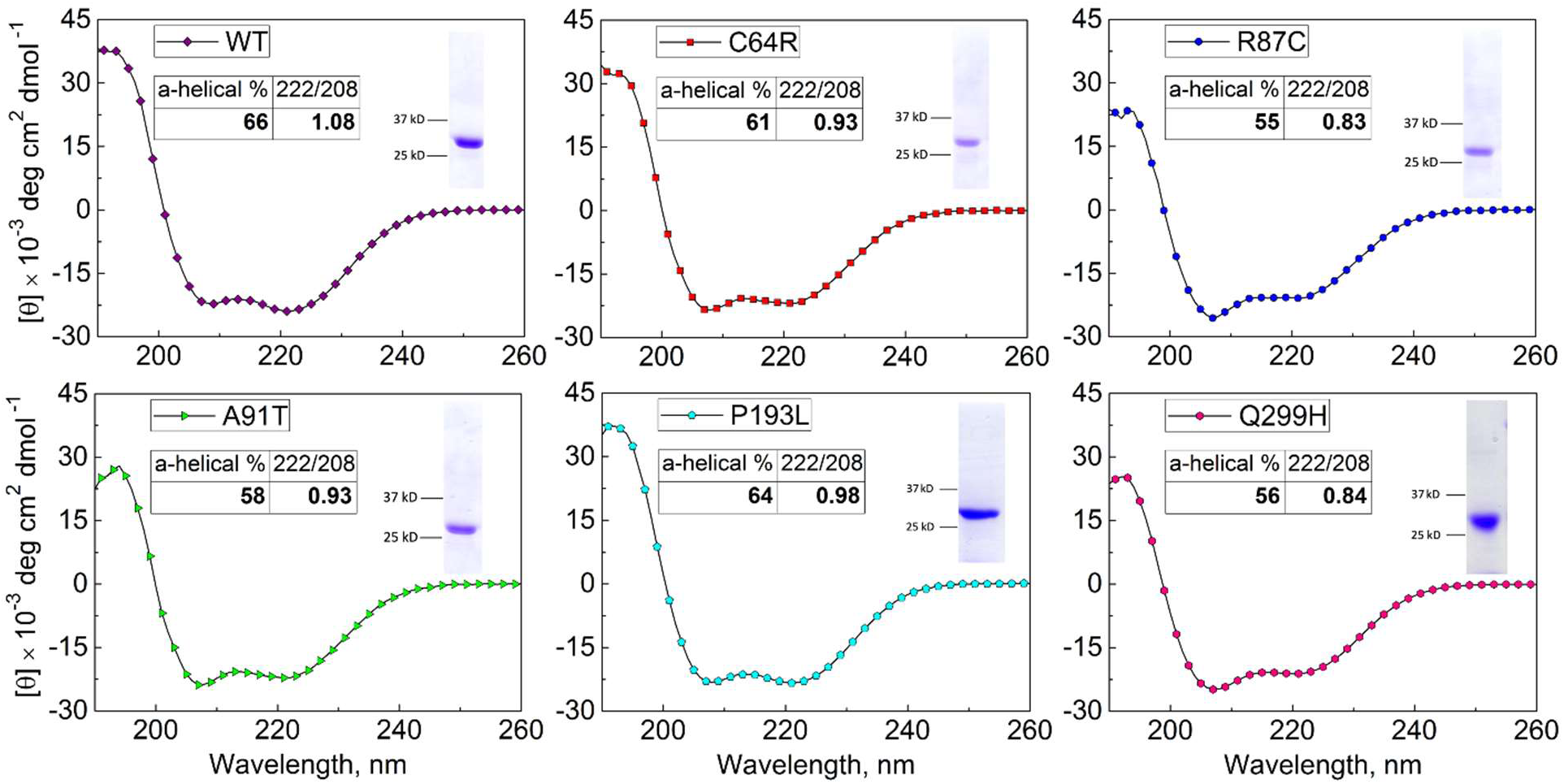
CD spectra of pZIP4-ECD and the variants. The α-helical content was estimated using the K2D3 server. 222/208 indicates the ratio of θ_222_/θ_208_. The SDS-PAGEs of purified protein are shown in the insets.

### The AE-causing mutations affect thermal stability and heat denaturation-associated thermodynamic parameters of pZIP4-ECD

Next, we compared the thermal stabilities of the wild type and the variants of pZIP4-ECD by gradually increasing temperature and recording changes of ellipticity at 222 nm in CD spectrum as an indicator of folding status. For the wild type protein, increasing temperature from 20 °C to 95 °C was accompanied with a decrease of θ_222_ and the plateau was reached at the end (Figure 6) when the protein was mostly denatured with α-helical content dropping from 66% to 20% (Figure S3). Although the ECD has two structurally distinct domains (HRD and PCD) with the PCD dimerizing through a large hydrophobic interface, the presence of only one transition in the heat denaturation curve suggests that the physical interactions between two domains may synchronize their unfolding processes. We also noticed that, when the heat-denatured protein was allowed to refold at room temperature, majority of the refolded protein is still a homodimer in solution as indicated in size exclusion chromatography, even though the α-helical content is decreased by 10% and the θ_222_/θ_208_ ratio is reduced to 0.82 (Figure S3). This result suggests that dimerization is either preserved at 95 °C or re-formed during protein refolding. In either scenario (or a mixed process), this result indicates that pZIP4-ECD has a strong tendency to keep and/or form a dimer, which is consistent with the proposed ECD function of facilitating dimerization of ZIP4 (19). By plotting normalized Δθ_222_ against temperature, the melting temperature (T_m_), at which the unfolding process reaches 50% during heat denaturation, can be estimated. When compared to the T_m_ of the wild type protein (59.5 °C), all the variants except for Q299H exhibited lowered T_m_s, ranging from 56.2 °C to 58.1 °C (Figure 6). The unexpected high T_m_ for the Q299H variant (61.4 °C) is accompanied with a more flattened heat denaturation curve when compared to the sigmoid curves of the wild type or the other variants, which is implicative of impaired cooperativity and interaction network of the structural elements of this variant.

**Figure 6.**
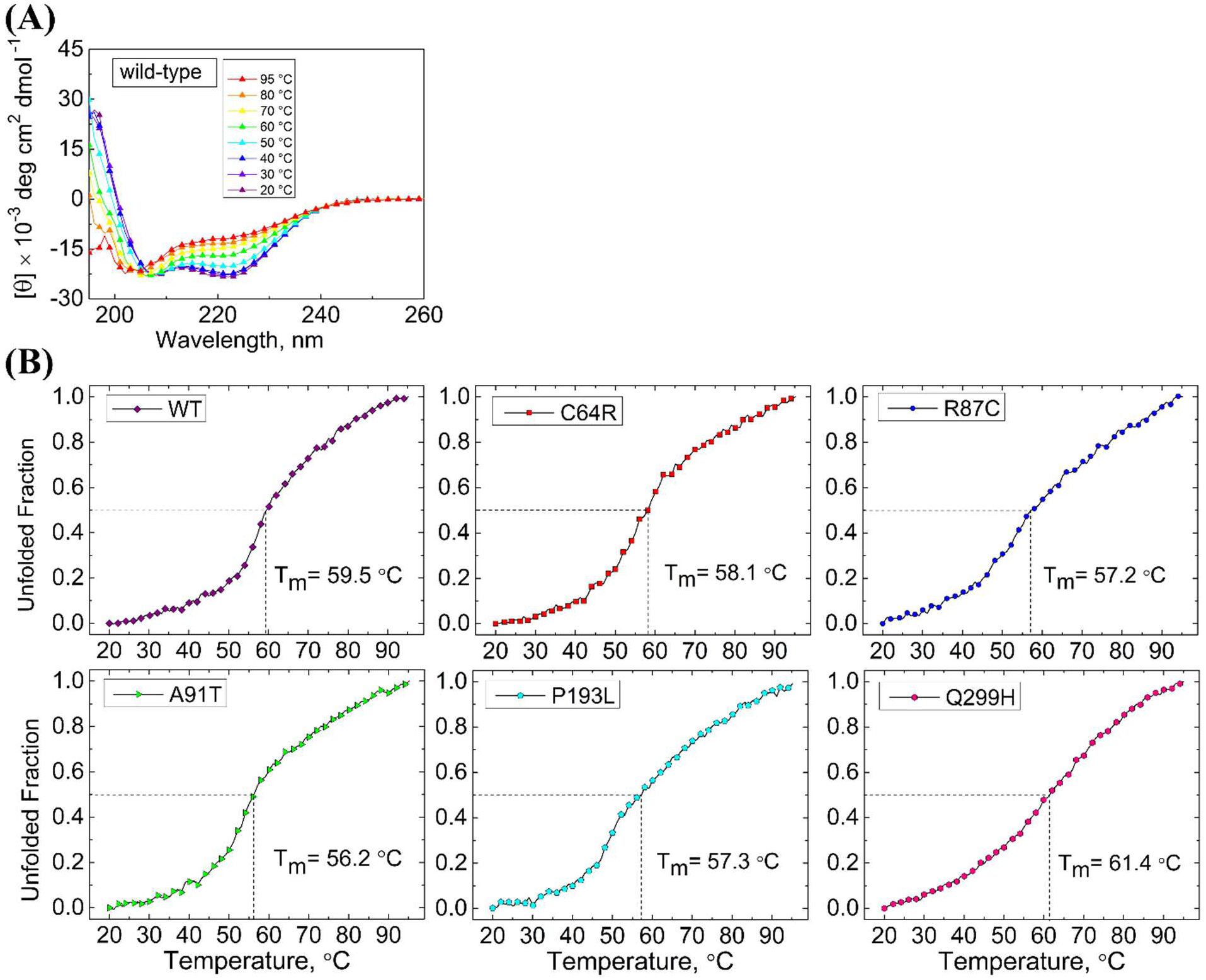
Heat denaturation of pZIP4-ECD and the variants. (A) CD spectra of wild type pZIP4-ECD at different temperatures. At 95 °C, the α-helical content was estimated to be 20% and further increasing temperature did not reduce helical content. (B) Heat denaturation curves of the wild type pZIP4-ECD and the variants. The melting temperature (Tm) was estimated on the denaturation curve where the unfolded fraction is 0.5.

Taken together, the CD spectra and the heat denaturation experiments of pZIP4-ECD variants revealed substantially changed biophysical properties, indicating that the substitutions in pZIP4-ECD, which are equivalent to the AE-causing mutations in hZIP4-ECD, lead to impaired protein folding and decreased thermal stability.

## Discussion

As the ZIP family plays a central role in transition metal homeostasis and cell signaling (18), LOF mutations of human ZIPs cause severe syndromes, including AE which is the first genetic disorder discovered to be associated with the ZIP family (5,6). So far, at least seven AE-causing mutations have been reported and some of them have been biochemically studied on mZIP4, a close homolog of hZIP4. Most of the missense mutations in the TMD of mZIP4 led to lower expression level, reduced cell surface expression and largely diminished zinc transport activity (7). Mapping the mutations on the structural model of hZIP4 generated using a bacterial ZIP structure has suggested that the involved residues appear to play structural roles and therefore the AE-causing mutations in the TMD likely affect protein folding and/or stability (35), which would eventually lead to degradation by quality control mechanism in the ER (31). The only characterized mutation in the ECD of mZIP4 is P200L, which is equivalent to P200L in hZIP4 (7). The same mutation in hZIP4 was also reported in a recent study (23). In this work, we characterized seven confirmed AE-causing mutations in the ECD (including P200L) and found that all the variants exhibited similar behavior when expressed in HEK293T cells – they were able to be expressed at similar (or modestly reduced) levels as the wild type hZIP4 but barely detected at cell surface, which accounts for the total loss of zinc transport activity. The fact that the variants are immaturely glycosylated and aberrantly trapped in the ER strongly indicates intracellular mistrafficking. By biophysically examining the effects of the corresponding mutations in pZIP4-ECD, we demonstrated that the mutations cause defects in folding and stability, which provides a causal explanation for mistrafficking and dysfunction of the AE-associated hZIP4 variants.

Protein misfolding and structural defects are known pathogenic mechanisms for dysfunction of membrane proteins caused by disease-linked mutations. A well-documented case is the autosomal recessive disorder cystic fibrosis (CF) which is caused LOF mutations of a chloride ion channel, cystic fibrosis transmembrane conductance regulator (CFTR). Notably, the most prevalent mutation (>90%) associated with CFTR is ΔF508, for which the residue F508 within the cytosolic nucleotide binding domain 1 (NBD1) is missed (36). The ΔF508 variant exhibited drastically reduced transport activity, immature glycosylation, reduced surface expression and accelerated clearance from plasma membrane (37). The atomic resolution of cryo-EM structures of CFTR have shown that F508 is buried in a hydrophobic core (38,39) which mediates physical interactions of the NBD1 and the TMD, and biochemical/biophysical studies have indicated that the variant has defects in folding and structural compactness (37). Remarkably, the AE-linked variants with single residue substituted in the ECD showed a similar phenotype as the ΔF508 CFTR variant. As discussed in our previous report and also shown in Figure 1, the AE-causing mutations occur at the conserved residues in structured regions where the involved residues play structural roles in formation of disulfide bonds (C62R, C309Y), salt bridge (R95C), hydrogen bonds (N106K and Q303H), and hydrophobic cores (A99T, P200L). With no surprise, the biophysical studies on pZIP4-ECD indicated that the tested five variants (C64R, R87C, A91T, P193L, and Q299H) have shown defects in folding and structural integrity, which are manifested by reduced α-helical content and the θ_222_/θ_208_ ratio (for all the variants), and decreased thermal stability (with the exception of the Q299H variant) (Figures 5 and 6). The extremely low yield for the D98K and C305Y variants already implies severe defects in folding and/or stability.

We notice the discrepancy between our results and other reports on the P200L variant. It was shown that the P200L variant of mZIP4 was expressed in HEK293T cells with a cell surface level comparable to that of the wild type protein and a 70% decrease in zinc transport activity (7). In contrast, the P200L variant of hZIP4 showed little surface expression and no detectable activity in this work (Figures 2 & 3). As already mentioned in the mZIP4 study, nearly all the wild type mZIP4 was maturely glycosylated, whereas only 50%-70% of the P200L variant did so, suggesting that a significant amount of the variant was not properly processed. Notably, even for the wild type hZIP4, the percentage of maturely glycosylated protein is just about 50% (Figures 2 and 3). It is unclear why mZIP4 is more efficiently processed and traffic in a human cell line than hZIP4. One possibility would be that mZIP4 is intrinsically more stable than hZIP4 so that it has better chance to fold correctly in the ER. It would also explain why the P200L variant of mZIP4 can still be presented at cell surface despite of folding defect indicated by immature glycosylation. While this work was being conducted, we noticed a study reporting that the P200L variant of hZIP4 expressed in HEK293T cells was shown to have a normal zinc transport activity which was measured by using a small molecule zinc-responsive fluorescence dye (23). It is unclear whether this discrepancy is due to different approaches used for tracing zinc transport. Radioactive ^65^Zn based transport assay has been used to study transport kinetics of a variety of ZIPs, including hZIP4 (7,19,28,29,35,40–44). Consistent with loss of zinc transport activity, the P200L variant of hZIP4 is shown to be barely expressed at cell surface. We detected cell surface expression levels of hZIP4 and its variants using two approaches following the previous mZIP4 study (7) – Western blot for detecting cell surface bound anti-HA antibody and immunofluorescence imaging for visualizing fluorescence-labeled anti-HA antibody associated at cell surface. As shown in Figures 3A and 3B, the results of both approaches consistently indicated that the AE-associated variants, including the P200L variant, had much lower levels of cell surface expression than the wild type hZIP4. Co-localization experiment further revealed that the P200L variant is largely retained in the ER (Figure 4), which explains the lack of zinc transport activity of this variant.

After the discovery of the causal link between ZIP4 mutations and AE, disease-causing mutations have been reported in other human ZIPs. The LOF mutations of ZIP13 cause the spondylocheirodysplastic form of Ehlers-Danlos syndrome (SCD-EDS) (42,45,46), an autosomal recessive disorder characteristic of defects of connective tissues, bones, and teeth. It has been demonstrated that both the missense mutation G64D and the deletion mutation ΔFLA (on the third transmembrane helix) drastically reduced stability and protein expression level in cells due to accelerated clearance through the ubiquitination dependent proteasome degradation pathway (45). Recently, inherited LOF mutations of ZIP8 and ZIP14, which are close homologs with broad substrate spectrum covering from the beneficial trace elements (zinc, iron, manganese) to the toxic cadmium, have been linked with severe failure of systemic manganese homeostasis (47,48). The disease-linked ZIP8 variants fail to transport Mn^2+^ because of abrogated cell surface expression and being aberrantly retained in the ER (49). Notably, these ZIP8 variants harbor at least one substitution in the ECD, highlighting the importance of the ECD in ZIP8 intracellular trafficking. In this work and also in our previous report, we demonstrate that the mutations on the conserved residues in the ECD significantly affected ZIP4 intracellular trafficking (19). Although the exact role of the ECD is still not fully clarified, these studies have suggested that correct folding and structural integrity of the ECD are required for proper intracellular trafficking of the ZIPs. Whether the ECD plays any active roles in folding, processing or trafficking of mammalian ZIPs would be interesting to investigate later.

In summary, seven confirmed AE-causing mutations were functionally and biophysically characterized in this work, and the results indicate that the mutations in the ECD lead to protein mistrafficking because of impaired folding, reduced thermal stability and undermined structural integrity, which accounts for the abolished zinc transport activity. Given that AE and CF share a similar molecular pathogenic mechanism and that small molecule chemical chaperones have been successfully applied for CF treatment (50), developing AE-specific chemical chaperones may represent an alternative strategy to treat this rare disease. After all, life-time and high-dose zinc administration may cause imbalance of other trace elements, such as copper deficiency (51,52).

## Experimental procedures

### Genes, plasmids and mutagenesis

The plasmids harboring hZIP4 (GenBank code: BC062625) or pZIP4-ECD (gene ID: ELK11751, residue 36-322) are the same as reported (19). All the site-directed mutations were conducted using QuikChange mutagenesis kit (Agilent, Cat# 600250) and verified by DNA sequencing.

### Cell culture, transfection and Western blot

Human embryonic kidney cells (HEK293T, ATCC, Cat #CRL-3216) were cultured in Dulbecco’s modified eagle medium (DMEM, Thermo Fisher Scientific, Invitrogen, Cat#11965092) supplemented with 10 % (v/v) fetal bovine serum (FBS, Thermo Fisher Scientific, Invitrogen, Cat#10082147) and Antibiotic-Antimycotic solution (Thermo Fisher Scientific, Invitrogen, Cat# 15240062) at 5% CO_2_ and 37 °C. Cells were seeded on poly-D-lysine (Corning, Cat# 354210) coated 24-wells trays for 16 hours in the basal medium and transfected with 0.5-0.8 µg DNA/well by lipofectamine 2000 (Thermo Fisher Scientific, Invitrogen, Cat# 11668019) in OPTI-MEM medium.

For Western blot, all the samples were heated at 96 °C for 6 min before loading on SDS-PAGE gel. The protein bands were transferred to PVDF membranes (Millipore, Cat# PVH00010). After being blocked with 5% non-fat dry milk, the membranes were incubated with anti-HA antibody (Thermo Fisher Scientific, Cat#26183) at 4 °C overnight. Bound primary antibodies were detected with HRP-conjugated goat anti-mouse immunoglobulin-G at 1:5,000 (Cell Signaling Technoloy, Cat# 7076S) or goat anti-rabbit immunoglobulin-G at 1:2,500 (Cell Signaling Technology, Cat# 7074S) by chemiluminescence (VWR, Cat# RPN2232). The blots were taken using a Bio-Rad ChemiDoc™ Imaging System.

### Zinc transport assay

Twenty hours post-transfection, cells were washed with the wash buffer (10 mM HEPES, 142 mM NaCl, 5 mM KCl, 10 mM glucose, pH 7.3) followed by incubation with 10 μM ZnCl_2_ (containing 30% 65ZnCl_2_) in Chelex-treated 10% FBS/DMEM media for 20 mins at 37 °C. Then, plates were transferred on ice and zinc uptake was stopped by addition of precooled 1 mM EDTA containing wash buffer. Cells were centrifuged at 3500 rpm and the supernatant was discarded. Cells were washed twice before lysis with 0.5% Triton X-100. Packard Cobra Auto-Gamma counter was used to detect radioactivity. The same procedure was used for activity assay of the P200L variant at designated zinc concentrations. The cells transfected with the empty vector were treated in the same way.

### hZIP4-HA surface expression detection

hZIP4-HA expressed at the plasma membrane was indicated by the surface bound anti-HA antibodies recognizing the C-terminal HA tag of hZIP4. After 24h transfection, cells were washed twice with DPBS on ice and then fixed for 10 min in 4% formaldehyde at room temperature. Cells were then washed three times in DPBS (5 minutes each wash) and incubated with 2.5 µg/ml anti-HA antibody diluted with 5% BSA in DPBS for 1.5 h at room temperature. Cells were washed seven times with DPBS to remove unbound antibodies and then lysed in SDS–PAGE loading buffer. The anti-HA antibody bound to the surface hZIP4-HA in cell lysates were detected in Western blot with HRP-conjugated goat anti-mouse immunoglobulin-G (1:5000). As loading control, β-actin levels were detected using an anti-β-actin antibody (1:2,500).

### Immunofluorescence imaging

HEK293T cells were grown in 24-well trays for 16 h on sterile glass coverslips and transfected with plasmids harboring the genes of hZIP4 or its variants using lipofectamine 2000. To visualize cell surface expressed hZIP4 or its variants, cells were washed briefly by DPBS after 24h transfection and then fixed for 10 min at room temperature using 4% formaldehyde. The cells were washed in DPBS (5 minutes each wash) and incubated with 2.5 µg/ml an FITC-labeled anti-HA antibody (Sigma, Product # H7411) diluted with 5% BSA in DPBS for 1.5 h at room temperature. After five washes with DPBS, coverslips were mounted on slides with fluoroshield mounting medium with DAPI (Abcam, Cat# ab104139). Images were taken with a 40X objective using a Spectral-based Olympus FluoView 1000 confocal laser scanning microscope (CLSM). To detect intracellular localization of hZIP4 variants and calreticulin (an ER marker), after fixation by using formaldehyde, cells were permeabilized and blocked for 1h with DPBS containing 5% goat serum (Cell Signaling Technology, Cat# 5425S) and 0.1% Triton X-100 and then incubated with anti-HA antibodies at 1:500 (Thermo Fisher Scientific, Cat#26183) and/or anti-calreticulin antibodies at 1:300 (Thermo Fisher Scientific, Cat#PA3-900) at 4 °C for overnight. After three washes with DPBS (5-minute incubation for each wash), cells were incubated with Alexa-568 goat anti-mouse antibodies at 1:500 (Thermo Fisher Scientific, Cat# A-11004) and Alexa-488 anti-rabbit antibodies at 1:500 (Thermo Fisher Scientific, Cat# A-27034). After 3 washes with DPBS, coverslips were mounted on slides with fluoroshield mounting medium with DAPI (Abcam, Cat# ab104139). Images were taken with a 100X objective.

### Expression and purification of pZIP4-ECD and the variants

The wild type pZIP4–ECD and the variants were expressed as previously described (19). In brief, the E.coli cells of Origami™ B(DE3) pLysS (Novagen) transformed with the pLW01 vector were grown at 37 °C in lysogeny broth medium until OD_600_ ~ 0.6, and expression was induced by 0.2 mM IPTG before the cells were transferred to 16 °C for overnight growth. After cell lysis by sonication, the His-tagged proteins were purified using a nickel-nitrilotriacetic acid column. After removal of N-terminal His_6_-tag by thrombin, the proteins were dialyzed against a Tris-HCl buffer (pH 8.0) containing 10 mM EDTA, subjected to ion-exchange chromatography (Mono-Q, GE Healthcare), and then polished by size-exclusion chromatography (Superdex 200 10/300 GL, GE Healthcare) in a buffer containing 10 mM HEPES, pH 7.3 and 100 mM NaCl.

### Circular dichroism experiments

To prepare the samples for CD experiments, Thermo Scientific Slide-A-Lyzer MINI Dialysis Device was used for changing the sample buffer to 10 mM phosphate pH 7.3 over the course of a two-day dialysis at 4 °C with buffer change for three times. Sample concentrations were measured by using a NanoDrop ND-1000 Spectrophotometer and then adjusted to proper concentration using the dialysis buffer.

The circular dichroism (CD) spectrum of 8 μM pZIP4–ECD (or the variants) in 300 μl of 10 mM phosphate buffer (pH 7.3) were recorded in Hellma^®^ 1 mm QS cuvette sealed with PTFE stopper by using a JASCO-815 CD Spectrometer. Wavelength range was set between 190-260 nm, and data was collected at 1 nm wavelength increments with 1.5 s averaging time per wavelength point. Three scans were recorded and averaged. The K2D3 server (http://cbdm-01.zdv.uni-mainz.de/~andrade/k2d3/n) was used to estimate secondary structure components of the proteins. For thermal stability analysis of wild type protein, CD spectra were recorded in the range of 195-260 nm and starting temperature was set at 20 °C. Temperature was increased in 10 °C increments and waiting time was set to 5 minutes at each designated temperature before scanning. Three scans were recorded and averaged. After reaching 95 °C, temperature was cooled down to 20 °C and waited for 10 minutes before scanning. Three scans were recorded and averaged. For thermal stability study, ellipticities at 222 nm were recorded while the temperature increased from 20 °C to 95 °C. Rate of temperature increase was 3 °C/min with a temperature tolerance of 0.15 °C. The average of three repeats was used in data processing and analysis. Unfolded fraction at temperature T (F_unfolded,T_) was calculated as below:

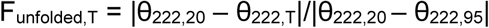

where θ_222,20_, θ_222,95_ and θ_222_,T are the readings of θ_222_ at 20 °C, 95 °C and temperature T, respectively. Tm was determined from the heat denaturation curve at which F_unfolded_ = 0.5.

## Acknowledgments

We thank Dr. Jin He at Department of Biochemistry and Molecular Biology for access to fluorescence microscopy for initial sample inspection. We thank Dr. Melinda Frame at Center for Advanced Microscopy for providing assistance in Confocal experiments.

## Author contributions

J.H. conceived the project and designed the experiments; E.K., C.Z. and D.S. conducted experiments; E.K., C.Z. and J.H. analyzed the data and wrote the manuscript.

---

**Figure S1.**
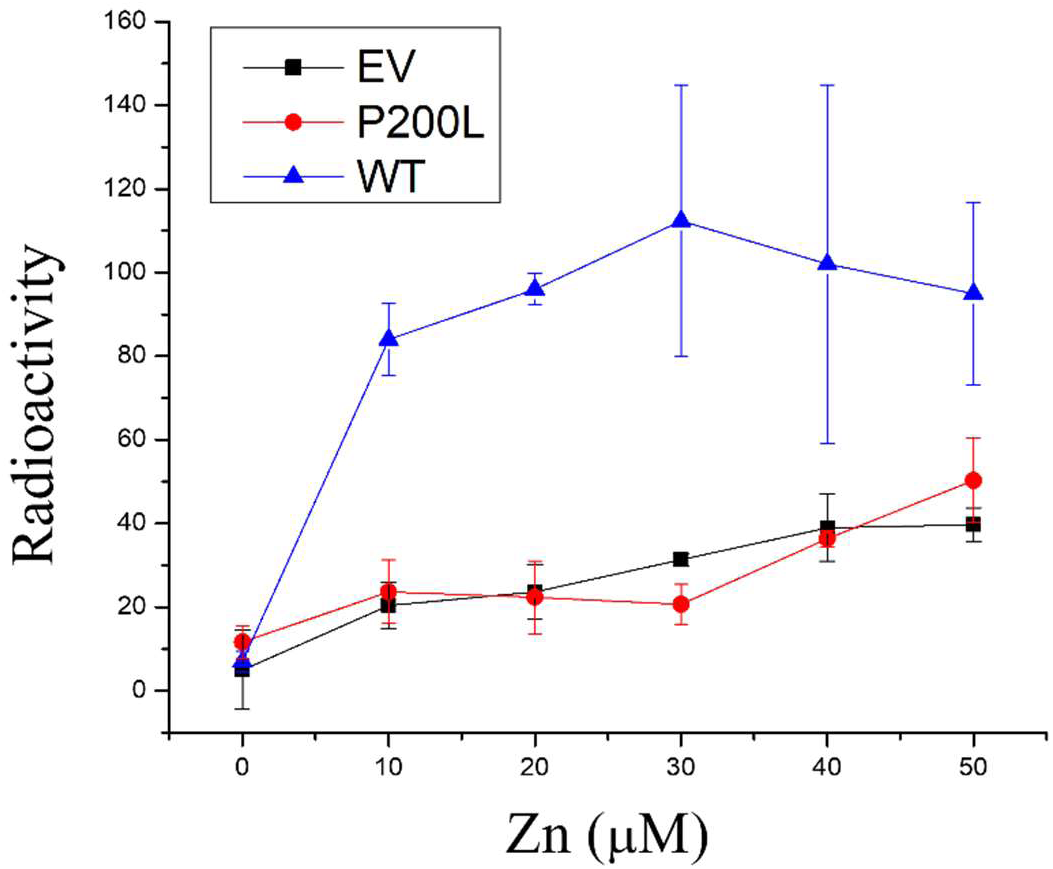
Zinc transport activity measurement at indicated zinc concentrations. Three technical repeats were averaged at each data point. The error bars indicate 1± standard deviation.

**Figure S2.**
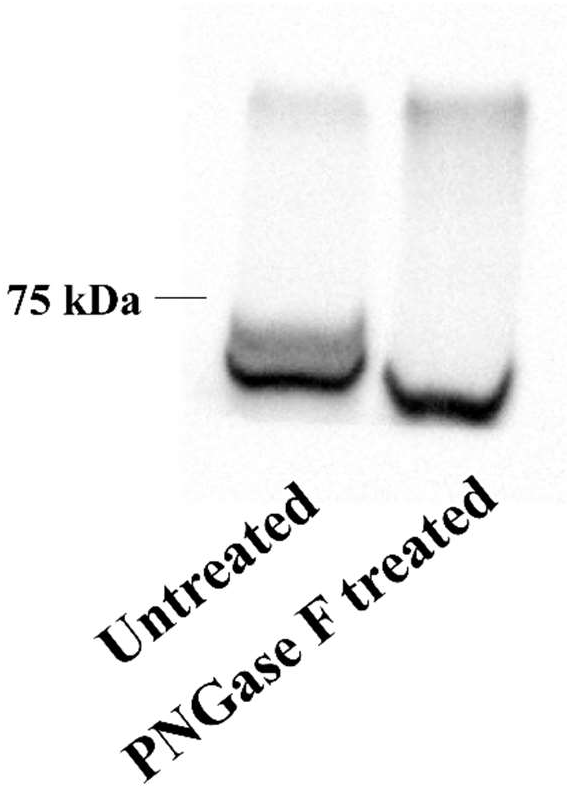
Processing of the wild type hZIP4 by PNGase F. PNGase F was directly added to the SDS-PAGE sample.

**Figure S3.**
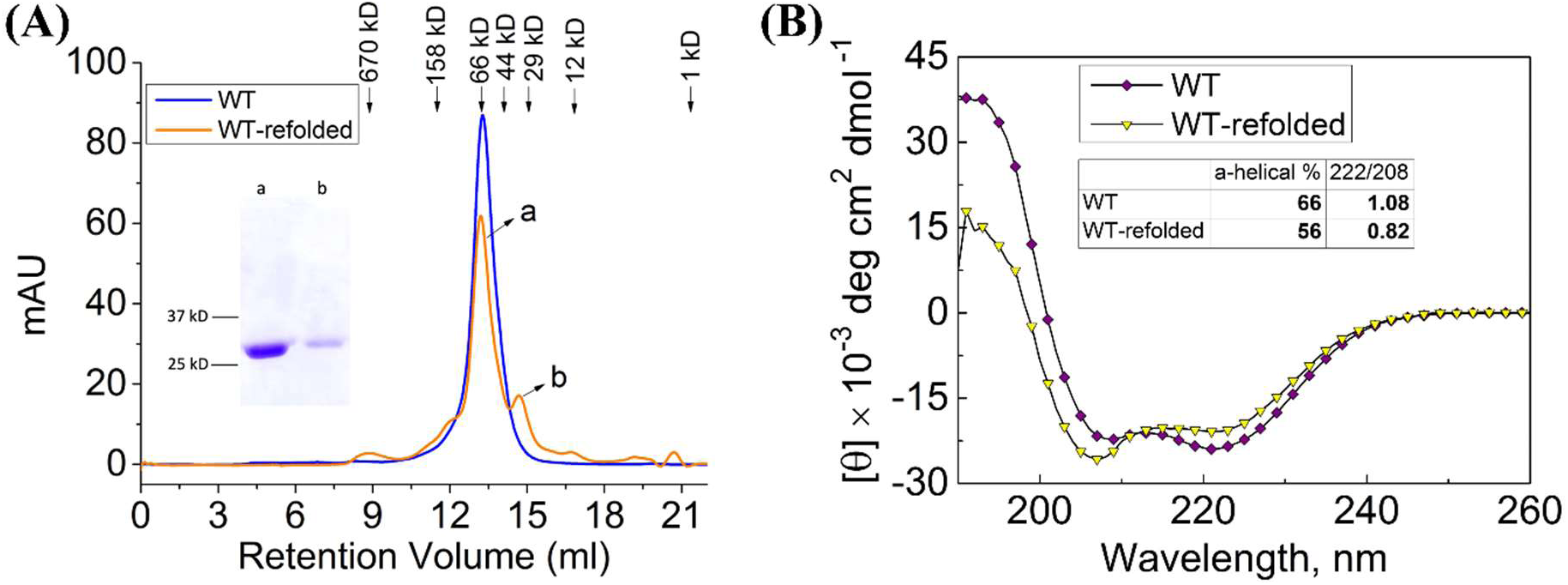
Characterization of refolded wild type pZIP4-ECD. (A) Size exclusion chromatography of the native and the refolded proteins. The elution volumes of the standard proteins are indicated by arrows. Note that the refolded protein has a major peak (peak a, dimer) and a minor peak (peak b, monomer). (B) CD spectra of the native and the refolded proteins. The refolded protein has lower α-helical content and smaller θ_222_/θ_208_ ratio than the wild type protein.

